# Identification of clinically approved small molecules that inhibit growth and promote surface remodeling in the African trypanosome

**DOI:** 10.1101/776658

**Authors:** Madison Elle Walsh, Eleanor Mary Naudzius, Savanah Jessica Diaz, Theodore William Wismar, Mikhail Martchenko, Danae Schulz

## Abstract

*Trypanosoma brucei* are unicellular parasites endemic to Sub-Saharan Africa that cause fatal disease in humans and animals. Infection with these parasites is caused by the bite of the tsetse fly vector, and parasites living extracellularly in the blood of infected animals evade the host immune system through antigenic variation. Existing drugs for Human and Animal African Trypanosomiasis are difficult to administer and can have serious side effects. Resistance to some drugs is also increasing, creating an urgent need for alternative trypanosomiasis therapeutics. In addition to identifying drugs that inhibit trypanosome growth, we wish to identify small molecules that can induce bloodstream form parasites to differentiate into forms adapted for the insect vector. These insect stage parasites do not vary proteins on their cell surface, making them vulnerable to the host immune system. To identify drugs that trigger differentiation of the parasite from bloodstream to insect stages, we engineered bloodstream reporter parasites that express Green Fluorescent Protein (GFP) following induction of the invariant insect-stage specific procyclin transcript. Using these bloodstream reporter strains in combination with high-throughput flow cytometry, we screened a library of 1,585 U.S. or foreign-approved drugs and identified eflornithine, spironolactone, and phenothiazine as small molecules that induce transcription of procylin. Both eflornithine and spironolactone also affect transcript levels for a subset of differentiation associated genes. We further identified 154 compounds that inhibit trypanosome growth. As all of these compounds have already undergone testing for human toxicity, they represent good candidates for repurposing as trypanosome therapeutics. Finally, this study is proof of principle that fluorescent reporters are a useful tool for small molecule or genetic screens aimed at identifying molecules or processes that initiate remodeling of the parasite surface during life cycle stage transitions.

**Author Summary:** African trypanosomes are unicellular parasites that infect humans and animals, causing a fatal disease known as sleeping sickness in humans and nagana in cattle. These diseases impose a severe economic burden for people living in Sub-Saharan Africa, where parasites are transmitted to humans and animals through the bite of the tsetse fly. Parasites living outside cells in humans and animals are attacked by the antibodies of the host immune system, but they can evade this attack by varying the proteins on their cell surface. In contrast, because flies do not have an antibody-mediated immune response, parasites living in flies do not vary the proteins on their cell surface. In this study, we performed a small molecule screen to identify compounds that might force bloodstream parasites to move forward in their life cycle to become more similar to parasites living in flies, causing them to express a protein on their cell surface that does not vary. This invariant protein on the surface of bloodstream parasites would make bloodstream parasites vulnerable to the host antibodies. We found 3 compounds that increased RNA levels for an invariant insect-stage surface protein and 154 compounds that inhibit parasite growth. We hope these compounds might have potential as novel trypanosomiasis therapeutics.

## Introduction

*Trypanosoma brucei* is a unicellular protozoan parasite that causes both Human and Animal African Trypanosomiasis (also known as sleeping sickness in humans and nagana in livestock, respectively). These diseases affect both humans and ungulates, causing a severe human and economic burden in regions of Sub-Saharan Africa where they are endemic (1). Because trypanosomiasis primarily affects the rural poor, there is a paucity of efficacious drugs available. Moreover, all drugs in current use have severe side effects, some drugs used to treat late stage trypanosomiasis are highly toxic, and resistance to existing drugs is rising (2). Thus, new strategies to treat trypanosomiasis are urgently needed.

Trypanosomes live extracellularly within the mammalian host and untreated infections are almost invariably fatal. This is because *T. brucei* has evolved a number of mechanisms that are effective at evading the mammalian host antibody response. Many of these depend on the expression and recycling of the Variant Surface Glycoprotein (VSG) coat that densely covers the surface of bloodstream trypanosomes (3). There are thousands of antigenically distinct genetic variants of *VSG* in the genome (4), and by periodically switching the variant expressed on the parasite surface, the host antibody response can be effectively thwarted. This immune evasion allows the parasite to live long enough within the mammalian bloodstream to be transferred to the tsetse fly insect vector following a bloodmeal, where it transitions to the insect-specific procyclic form. The environment of the fly is quite different from that of the mammalian bloodstream, and the parasite adapts itself accordingly by changing its morphology and metabolism (5). Indeed, it’s been shown that thousands of genes show altered transcript levels between bloodstream and procyclic forms (6–8). A key event marking the transition from bloodstream to procyclic forms is the remodeling of parasite surface proteins to replace VSG with an invariant surface protein called procyclin, of which there are only four isoforms (9). The parasites must then make their way from the insect midgut to the salivary gland to be transferred back to a mammalian host.

Much progress has been made in elucidating the quorum sensing signaling pathways that are necessary to trigger parasite differentiation from the bloodstream form to the insect form via the stumpy intermediate (10–12). It is also clear that RNA Binding Proteins (RBPs) play an important role in maintaining life cycle stage-specific proteins and triggering differentiation events (13–16). However, the gene regulatory mechanisms by which VSG transcription is halted and procyclin transcription is initiated and/or stabilized remain poorly characterized. In order to learn more about the molecular mechanisms that initiate transcription of the *EP1* gene that codes for procyclin, we designed a fluorescent reporter system to allow us to easily detect transcript levels of *EP1* using flow cytometry.

The advantages of this *EP1* reporter system are three-fold. First, it allows us to perform high-throughput flow cytometric-based small molecule screens to identify compounds that result in the expression of the invariant procyclin protein in bloodstream form parasites. The expression of an invariant protein on the surface of bloodstream cells provides a ‘handle’ that the mammalian immune system can use to eliminate parasites whose surface is usually a moving target due to the ever-shifting repertoire of VSGs. Thus, we can identify compounds that make the bloodstream parasite vulnerable to the host immune system, potentially identifying urgently needed new trypanosomiasis therapeutics. As an example, we previously showed that the small molecule bromodomain inhibitor I-BET151 has this effect in bloodstream form cells, resulting in the expression of procyclin on the surface and decreased virulence in a mouse model of infection (17).

The second advantage of such a reporter system is that the flow cytometry-based screen allows simultaneous screening for small molecules that are trypanocidal or that inhibit parasite growth, further increasing the potential to identify new therapeutics. Finally, our long-term goal is to take advantage of this reporter system to perform genetic screens, allowing us to identify the key players responsible for initiating and/or stabilizing procyclin transcripts during differentiation from the bloodstream form to the procyclic form. This will allow us to unravel the missing link between environmental signals for differentiation that are received and transduced at the parasite surface and the downstream effect on transcript levels that ultimately allow the parasite to adapt from the bloodstream environment to the insect environment.

Here we present proof of principle that our *EP1* fluorescent reporter can be used for flow cytometry-based screening using an autosampler. This allows for automated 96 well plate sampling without the need for a human operator, resulting in more high-throughput flow cytometry-based drug library screens. First, we showed that fluorescence levels in the reporter line faithfully mimic expected levels of *EP1* transcript in bloodstream, procyclic, and differentiating parasites. Second, we conducted a proof of principle small molecule screen using the Johns Hopkins University Clinical Compound Library and identified several compounds that increase *EP1* transcript levels in bloodstream form parasites. Finally, we were able to identify a ∼ 150 small molecules that inhibit parasite growth, many of which have not previously been reported.

## Methods

### Culture growth and strain details

We used the Lister 427 L224 strain of bloodstream *T. brucei* parasites with a dual BES marked line that expresses *VSG3* from BES7 (with Neo^R^ downstream of the promoter) and a Puro^R^ gene downstream of the BES2 promoter (expressing *VSG2*)(18). Bloodstream parasites were cultured in HMI-9 at 37°C with 5% CO_2_. Procyclic PF427 parasites were grown at 27°C in SDM79.

### Generation of the EP1/GFP reporter plasmid

DNA encoding GFP flanked by 314 bp immediately upstream of the *EP1* ORF and 279 bp immediately downstream of the *EP1* ORF was synthesized by Life Technology. Both upstream and downstream *EP1* fragments contain the entire 5’ and 3’ UTRs of *EP1*. The 5’UTR GFP 3’UTR fragment was amplified with PvuII and HindIII sites and cloned into the pyrFEKO Hygro^R^ plasmid using the same sites. We then amplified a 289bp homology region downstream of the *EP1* 3’UTR fragment with SpeI and XhoI, which was cloned downstream of the Hygro^R^ gene using the same sites. The *5’UTR/GFP/3’UTR* fusion fragment was amplified using Forward ggccatcagctgagtcaatagtgcattttg and Reverse ggccataagcttgaatgaaaaaaaatagaagtgaaa primers. The downstream homology region was amplified using Forward ggccatactagtctttgaatttggatcttaaaattattattg and Reverse ggccatctcgagcaacttcagctgcggggc primers. For transfection into our Hygro resistant procyclic strain, the construct was modified to replace the Hygro^R^ gene with a Puro^R^ gene.

### Strain construction

Transfection of bloodstream form parasites was performed using an AMAXA Nucleofector. Parasites were resuspended in 100µl T cell solution with 10µg linearized DNA and electroporated on the X001 setting. Following transfection, cells were grown in 5µg/ml Hygromycin (Invivogen). PF cells were transfected using the same procedure, but following transfection the cells were plated into conditioned media in 0.1µg/ml Puromycin (Invivogen). Correct integration of the construct in transfected cells was tested using PCR with a forward primer upstream of the 5’UTR of *EP1* (agtccgataggtatctcttattagtatag) and a reverse primer within the *GFP* gene (agaagtcgtgctgcttcatgtggt). Successful isolation of genomic DNA was confirmed using control primers that amplify the downstream *EP1* homology region: For ggccatactagtctttgaatttggatcttaaaattattattg and Rev ggccatctcgagcaacttcagctgcggggc.

### Differentiation

Bloodstream parasites were spun down and resuspended in differentiation media (19) supplemented with 6 mM cis-aconitate (Sigma-Aldrich A3412) at 27°C for three days.

### Sytox Orange Staining

Cells were resuspended in 5µM Sytox Orange (Fisher Scientific S11368) and incubated for 15 minutes at 37°C prior to flow cytometric analysis.

### Flow cytometry

All flow cytometry was done with a Novocyte 2000R from Acea biosciences. Procylin protein expression was measured following three to five days of drug treatment at 33 µM, after which parasites were stained for 10 minutes on ice with anti-EP1 (Cedarlane CLP001A) antibody. Cells were washed twice in HMI9 prior to analysis.

### Data analysis and statistical tests

#### Drug screen

A total of 1,585 drugs were tested from the Johns Hopkins University Clinical Compound Library, which had patented compounds removed prior to drug screening. Drug screening was conducted with the *EP1/GFP* parasites plated with 1,000 cells in 100 µL cultures and 33 µM of drug. After 3 days of growth, cells were screened for GFP fluorescence via flow cytometry. To test for autofluorescence of candidate hits, 300,000 cells/100µL culture were plated with 33 µM of each drug and immediately screened for fluorescence via flow cytometry. Each plate also contained three wells of negative control cells treated with DMSO and a well of positive control cells treated with 20µM I-BET151 (Sigma-Aldrich).

All samples with an average median GFP fluorescence intensity that was 1.5-fold or higher than the +DMSO control from the first two rounds of screening were selected for further screening. Drugs tested in triplicate were subjected to a two-sided, unpaired Student’s t-test. To identify drugs that inhibit trypanosome growth or those that are trypanocidal, we used flow cytometry to calculate total cell counts/10µl of volume. Control cells typically showed ∼ 35000 cells/10µl volume within the live gate, so we identified trypanocidal drugs as those that resulted in counts between 350-3500, 35-350, and <35 in the live gate.

#### Confirmation of drug hits

1 mL cultures of *EP1/GFP* reporter cells were treated with 33 µM of eflornithine (Sigma-Aldrich 1234249), phenothiazine (Sigma-Aldrich P14831), spironolactone (Sigma-Aldrich S3378), or vehicle control and grown for three days. The cells were then screened for GFP fluorescence via flow cytometry.

#### Growth curves

1 mL cultures of *EP1/GFP* reporter cells were plated at 100,000 cells/ml and treated with 33 µM of eflornithine (Sigma-Aldrich 1234249), phenothiazine (Sigma-Aldrich P14831), spironolactone (Sigma-Aldrich S3378), or vehicle control and counted daily by hemocytometer. Each day following counting, cells were diluted back to 100,000 cells/ml. For growth curve analysis by flow cytometry, 200 µl cultures of parasites were plated at 100,000 cells/ml and treated with 33 µM of phenothiazine (Sigma-Aldrich P14831), triprolidine hydrochloride (Sigma-Aldrich T6764), flunarizine hydrochloride (Sigma-Aldrich F8257), aprepitant (Sigma-Aldrich SML2215), bufexamac (Sigma-Aldrich B0760), clemastine fumarate salt (Sigma-Aldrich SML0445) or vehicle control. Parasites were stained daily with Sytox Orange and a fixed volume of cells was analyzed by flow cytometry. The number of live cells was calculated as the number of Sytox Orange negative cells falling within the live gate. Each day following flow analysis, cells were diluted back to 100,000 cells/ml according to the number of live cells counted.

#### IC_50_ Analysis

2000 parasites were plated in a 200µl volume of HMI9 (10000 cells/ml) at the indicated concentrations of each drug and grown for 48 hours at 37°C with 5% CO_2_. Parasites were then stained with Sytox Orange and the number of Sytox Orange negative parasites falling within the live gate was calculated using flow cytometry with a fixed volume of cells. Data were analyzed and fitted using GRMetric, an R package for calculation of dose response metrics based on growth rate inhibition (20). Growth rate values are calculated individually for each treatment using the formula

GR = 2 ^ (log_2_(cellcount/cellcount_control) / log_2_ (cellcount_control /cellcount_time0) – 1 The data are fitted with a sigmoidal curve

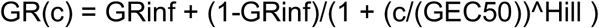

where GRinf = GR(count>inf), reflecting asymptotic drug efficacy. GEC50, the drug concentration at half-maximal effect, Hill, the Hill coefficient of the fitted GR curve reflecting how steep the dose-response curve is.

#### Cell cycle analysis

Cells were fixed using 1% formaldehyde in PBS for 5 minutes. Cells were then washed three times in PBS and then resuspended in a solution of 0.2 mg/ml RNAse and 0.05 mg/ml propidium iodide in PBS. Following a 2-hour incubation at 37°C, cells were analyzed by flow cytometry.

#### Quantitative PCR Analysis

For Q-PCR to quantify transcription, RNA was extracted from parasites treated with drug or vehicle control for 3 days using RNA Stat-60 (Tel-Test) following the manufacturer’s protocol and quantified on a NanoDrop2000c. 2.5μg of RNA was used to generate cDNA using the SuperScript IV VILO Master Mix (Fisher Scientific 11756050) according to the manufacturer’s protocol. cDNA was amplified using 2X Sybr green master mix (Life Technologies 4309155) and primers and quantified on an Eppendorf Realplex2 instrument. Primers used were as follows: *Tb427.10.10260 EP1*, tctgctcgctattcttctgttc, cctttgcctcccttagtaagac, *Tb927.6.510 GPEET* agtcggctagcaacgttatc, ttctggtccggtctcttct, *Tb927.9.11600, GIM5B*, ttgcgaggatgggtgatg, gggtttggagagggaagttaat, *Tb927.10.2010, HK1*, gtcagcacttactcccatcaa, acgacgcatcgtcaatatcc, *Tb927.10.5620, ALD*, gtctgaagctgttgttcgtttc, cacctcaggctccacaatag, *Tb927.10.10220, PAG2*, aggagatacgaggaatgagaca, tcttcaaacgcccggtaag, *Tb927.10.14140 PYK1*, gagaaggttggcacaaagga, tcacaccgtcgtcaacataaa, *GFP*, ctacaacagccacaaggtctat, ggtgttctgctggtagtg, *Tb927.10.9400, SF1*, ggtatggttcatcaggagttgg, cgttagcactggtatccttcag.

#### Mammalian cell growth assay

RAW264.7 mouse macrophage or C32 human melanoma cells were maintained in DMEM (Sigma-Aldrich) supplemented with 10% fetal bovine serum, as well as 100 μg/mL penicillin and 100 μg/mL streptomycin. 10,000 RAW264.7 cells per well were seeded in 96-well plates in 100 μl/well 24 hours before the assay. On the day of the assay, 0.75 or 1.5 μl of 3.3 mM JHCCL drugs were added to 150 μl of cell-containing media to achieve 16 or 33 μM of each compound per well, respectively. Cells were treated with drugs for 6 hours at 37 °C and 5% CO_2_. C32 cells were treated with drugs for 24 hours. Determination of cell viability by 3-(4,5-dimethylthiazol-2- yl)-2,5-diphenyltetrazolium bromide (MTT) assay was performed as described (21). Cell viability is shown as the percentage of surviving cells obtained relative to cells treated with DMSO as vehicle control (100%).

## Results

### Construction and validation of the *EP1/GFP* reporter cell line

In order to facilitate high-throughput screening for compounds that induce transcription of the *EP1* gene in bloodstream parasites, we integrated a construct containing *GFP* flanked by native EP1 5’ and 3’ UTRs at the endogenous *EP1* locus (Fig 1A). The construct was integrated in both bloodstream and procyclic parasites, and the second *EP1* allele was left intact. A PCR assay confirmed the correct integration of the reporter construct in both bloodstream and procyclic parasites (Fig 1A). We expect that if this reporter construct faithfully recapitulates native *EP1* expression, GFP levels should be relatively low in bloodstream form parasites and high in procyclic parasites. Fig 1B shows that GFP levels in *EP1/GFP* reporter cells are low in bloodstream parasites and high in procyclic parasites, as expected. We also expect that GFP expression in bloodstream *EP1/GFP* reporter parasites should increase when parasites are induced to differentiate to the procyclic form using cis-aconitate and incubation at 27°C, and we found that this was indeed the case (Fig 1C). Finally, we tested the effect of the bromodomain inhibitor I-BET151, which was previously shown to induce *EP1* expression in bloodstream parasites (17). Treatment of bloodstream *EP1/GFP* reporter cells with I-BET151 resulted in an increase in GFP expression after 3 days, and the effect was more pronounced after 5 days (Fig 1D). We conclude that *GFP* expression mirrors the expected expression of *EP1* in the *EP1/GFP* reporter line by all the metrics tested.

**Figure 1.**
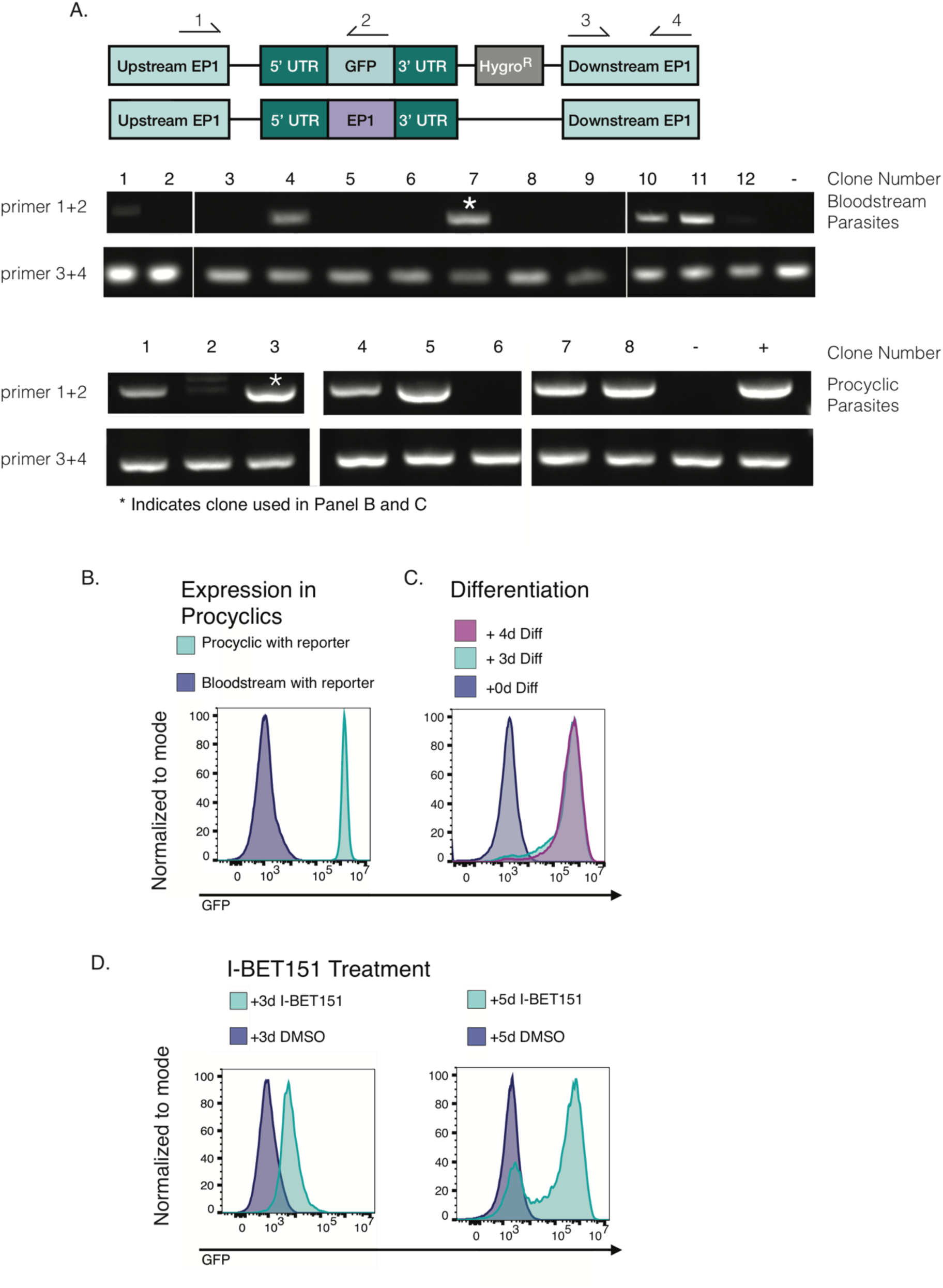
Validation of the *EP1/GFP* reporter construct. (A) (Top) Schematic showing the endogenous *EP1* locus following replacement of one allele with the *EP1/GFP* construct, wherein *GFP* is flanked by the native *EP1* 5’ and 3’ UTRs and marked with a Hygromycin resistance gene. The second *EP1* allele remains intact. Arrows indicate primers used to screen clones for correct integration of the reporter. Primers 1 and 2 were used to confirm integration, and primers 3 and 4 were used as a control for the presence of genomic DNA. (Middle) Gel electrophoresis of PCR products generated from 12 clones of bloodstream form parasites transfected with the *EP1/GFP* reporter construct. The parental line was used as a negative control (-). 4 clones showed correct integration by this assay, and (*) indicates the clone that was used for subsequent screening. Numbers on the left-hand side of the gel indicate the primer pair used. (Bottom) Same as the middle panel but for procyclic parasites transfected with the *EP1/GFP* reporter construct. The parental line was used as a negative control (-) and genomic DNA from a positively scoring bloodstream clone was used as a positive control (+). 6 clones scored positive and (*) indicates the clone used for flow cytometry in the panel below. (B) Flow cytometry plot for *GFP* expression in bloodstream and procyclic *EP1/GFP* reporter parasite clones. (C) Flow cytometry plot showing *GFP* expression in uninduced bloodstream *EP1/GFP* reporter parasites and in these same parasites induced to differentiate to procyclic forms using 6mM cis-aconitate and incubation at 27°C. (D) Flow cytometry plot showing *GFP* expression in bloodstream *EP1/GFP* parasites treated with the bromodomain inhibitor I-BET151 or vehicle control for 3 days (left) or 5 days (right).

### A proof of principle screen shows that the reporter strain can be used to identify small molecules that increase transcription of the procyclin gene *EP1*

To test whether the *EP1/GFP* reporter could be used to screen small molecules using high-throughput flow cytometry, we obtained the Johns Hopkins University Clinical Compound Library of 1,585 small molecule drugs. 1082 of these compounds are FDA approved while the remaining are foreign approved drugs. We plated 1000 bloodstream *EP1/GFP* reporter parasites in 100µl of media in a 96 well plate format. We then added 1µl of each drug to the well for a final concentration of 33µM. After 3 days of growth, we subjected each well to analysis by flow cytometry to measure the level of *EP1/GFP* expression. Each plate also contained parasites growing in DMSO to serve as a solvent control. This assay was performed in duplicate for each of the drugs. After the two initial assays, the median GFP fluorescence intensity was calculated for DMSO-treated parasites and drug treated parasites. Any drug that caused a 1.5- fold increase in average median GFP fluorescence intensity in drug-treated parasites compared to DMSO-treated control parasites was moved forward to the third round of screening. During this third round, in addition to the 3-day assay, drug-treated parasites were immediately subjected to analysis by flow cytometry following the addition of the drug to eliminate any drugs that were themselves fluorescent. After this third round, the median GFP fluorescence intensities for drug-treated parasites and solvent control-treated parasites were calculated again, and a Student’s t-test was performed to identify drugs that caused a significant increase in median GFP fluorescence intensity. For those drugs that induced a significant shift, we imposed a 2-fold median GFP fluorescence intensity cutoff (S1 Table). This left us with 9 candidate drugs, for which we ordered fresh stocks from Sigma-Aldrich. Parasites were again subjected to the 3-day flow cytometry assay, except this time they were grown in 1ml cultures. We were able to replicate the increase in *EP1/GFP* expression for Spirinolactone, Phenothiazine, and Eflornithine (Fig 2). Notably, all of our false positives for *EP1*/*GFP* expression inhibited cell growth (S1 Table). We hypothesize that the low cell counts produced noise in the data and that dying cells may have produced autofluorescence in the GFP channel. During follow-up experiments, the higher number of cells assayed by flow cytometry may have alleviated one or both of these problems.

**Figure 2.**
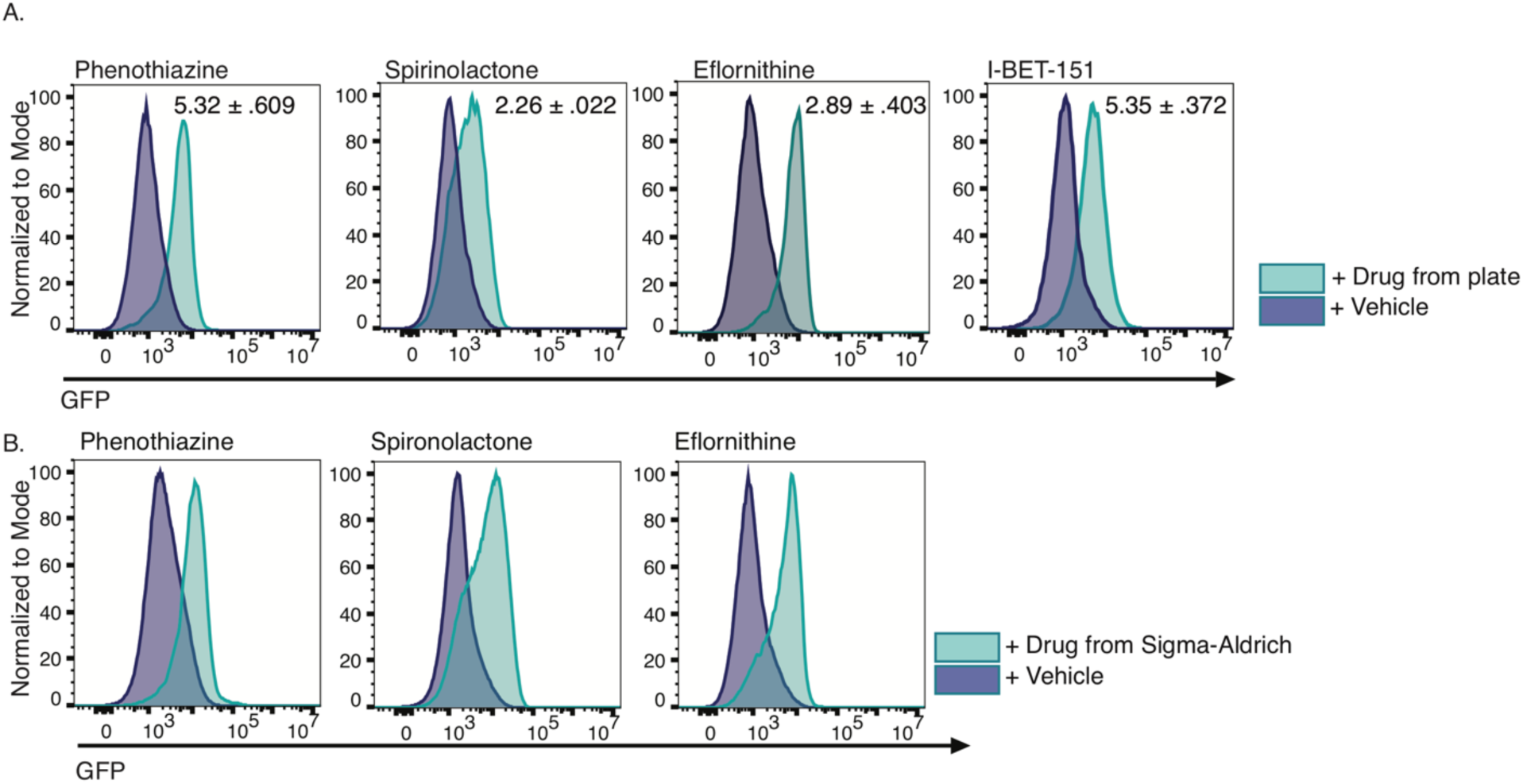
Three drugs from the small molecule library produced a reproducible shift in *EP1/GFP* expression. (A) Flow cytometry plots of *GFP* expression in *EP1/GFP* reporter parasites grown for 3 days in the presence of 33µM drug sourced from library plates or vehicle control. Inset indicates average fold change in median fluorescence intensity with standard deviation from 3 replicates of the experiment. (B) Flow cytometry plots of *GFP* expression in *EP1/GFP* reporter parasites grown for 3 days in the presence of fresh drug at 33µM sourced from Sigma-Aldrich.

Of the drugs we identified as increasing *EP1/GFP* expression, eflornithine treated bloodstream parasites have previously been reported to show some similarities to stumpy cells, although these stumpy-like forms are not competent to differentiate to procyclic forms (22). As stumpy forms have been shown to transcriptionally preadapt themselves to insect stage procyclic cells by upregulating procyclic specific transcripts (23), the increase in *EP1/GFP* expression in eflornithine treated parasites is consistent with these parasites sharing some characteristics of stumpy form parasites. Phenothiazine and its derivatives have previously been reported as anti-protozoal, strongly inhibiting cell growth in *T. brucei* (24,25). Lastly, to our knowledge, spironolactone has not been studied in the context of *T. brucei* infection, although it has been tested for its effects on myocardial protection in a model of Chagas disease (26). Spironolactone is used for the treatment hirsutism, as is eflornithine, which is one of the most important anti-trypanosomal drugs in current use (27).

### Phenothiazine causes severe growth defects and cell cycle abnormalities

We wished to determine if the observed increase in *EP1/GFP* expression for each candidate drug was a non-specific result of cell cycle arrest, so we tested the drugs that caused a measurable increase in *EP1/GFP* expression for their effect on parasite growth. While neither spironolactone nor eflornithine induced a profound effect on parasite growth at 33µM, phenothiazine caused a growth arrest in treated cells by 24h (Fig 3A). As expected, eflornithine did have effects on parasite growth at higher concentrations of 120µM (Fig 3A). To test the effect of each of these drugs on cell cycle, we performed propidium iodide staining and flow cytometry on parasites treated with each of the drugs for three days. In keeping with our observations of cell growth, phenothiazine treated parasites exhibited severe cell cycle abnormalities after 3 days of treatment (Fig 3B). We further tested phenothiazine treatment of parasites at earlier time points and found significant cell cycle abnormalities within 24h of treatment (Fig 3C). At 1 day the most pronounced effects were the accumulation of parasites in the G2 phase of the cell cycle, a decrease in S phase cells, and the appearance of multinucleated cells. At 2 days an accumulation of parasites lacking a nucleus (zoids) was apparent, and this effect was even more pronounced at 3 days (Fig 3D). The percent of parasites in each of the cell cycle stages was significantly different at all time points tested when compared to the DMSO treated controls. These results are in keeping with previously published reports that show that phenothiazine treated parasites exhibit severe growth defects through inhibition of trypanothione reductase (24,28). We conjecture that the effect of phenothiazine on *EP1/GFP* expression might be a non-specific stress response, while this does not appear to be the case for either spironolactone or eflornithine.

**Figure 3.**
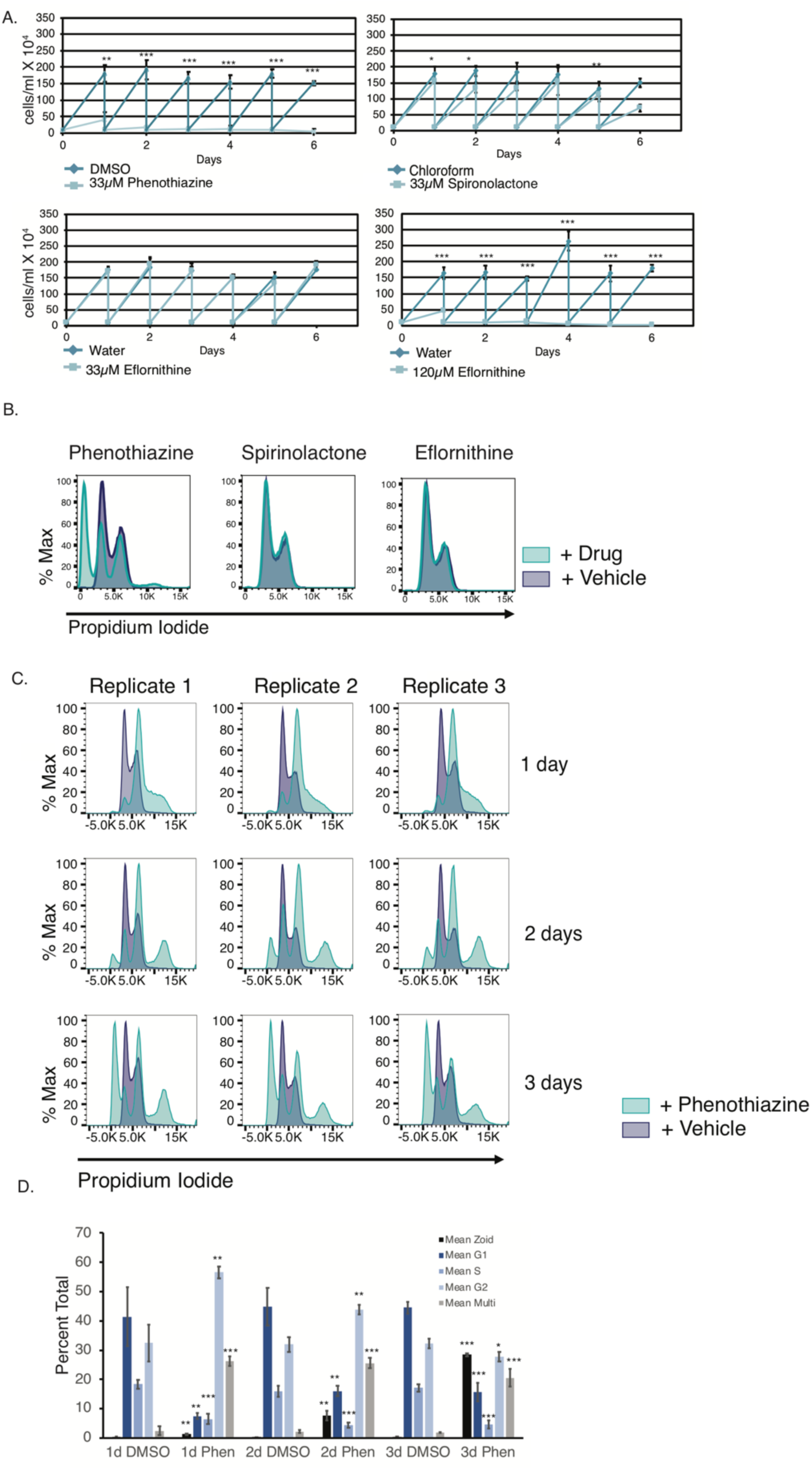
Phenothiazine-treated parasites are growth-arrested and show cell cycle defects. (A) Growth curves of bloodstream parasites grown in the presence of the indicated drug or vehicle control for 6 days. Parasites were counted daily on a hemacytometer and diluted to 100,000 cells/ml after counting. Error bars indicate standard deviation of 3 biological replicates. *, **, *** indicates p < 0.05, p < 0.01, and p < 0.001 respectively as measured by an unpaired two-sided Student’s t-test. (B) Flow cytometry plot using propidium iodide staining to assay cell cycle in bloodstream parasites treated with the 33µM of the indicated drug or vehicle control for 3 days. (C) Flow cytometry plot using propidium iodide staining to assay cell cycle in bloodstream parasites treated with the 33µM phenothiazine or vehicle control for the indicated number of days. (D) Quantification of the data collected in (C) showing the percent of cells in each phase of the cell cycle. *, **, *** indicates p < 0.05, p < 0.01, and p < 0.001 respectively as measured by an unpaired two-sided Student’s t-test.

### Surface protein expression of procyclin is not observed following treatment with drugs observed to increase *EP1/GFP* transcript levels

We tested whether the increase in *EP1/GFP* transcript levels corresponded to an increase in the corresponding procyclin protein on the parasite surface. Because our antibody to procyclin fluoresces in the same channel as GFP, we treated the parental strain for our reporter strain for 5 days with 30µM eflornithine, phenothiazine, and spironolactone, stained the parasites with anti-procyclin, and assayed procyclin protein expression by flow cytometry. I-BET151 was used a positive control as it has previously been shown to increase surface protein expression if procyclin (17). Previous experiments showed that the highest level of procyclin expression for I- BET151 was at 5 days so we used a similar treatment for our new drug candidates and replenished the drug every two days. None of the drugs produced measurable procyclin protein on the parasite surface after treatment (Fig 4). An increase in *EP1* transcription without a concomitant increase in protein expression has been observed before in the context of DNA replication mutants (29). While eflornithine is already in use as a trypanosome therapeutic and phenothiazine may have potential as a therapeutic based on its effect on cell growth, we conclude that spironolactone is not a promising drug candidate for trypanosomiasis since the effect on *EP1* transcript levels is not recapitulated at the level of protein (Fig 4, Fig 5). However, future studies beyond the scope of this work might aim to identify the protein target of spironolactone to try and ascertain whether this protein maintains low levels of *EP1* transcript in bloodstream cells.

**Figure 4.**
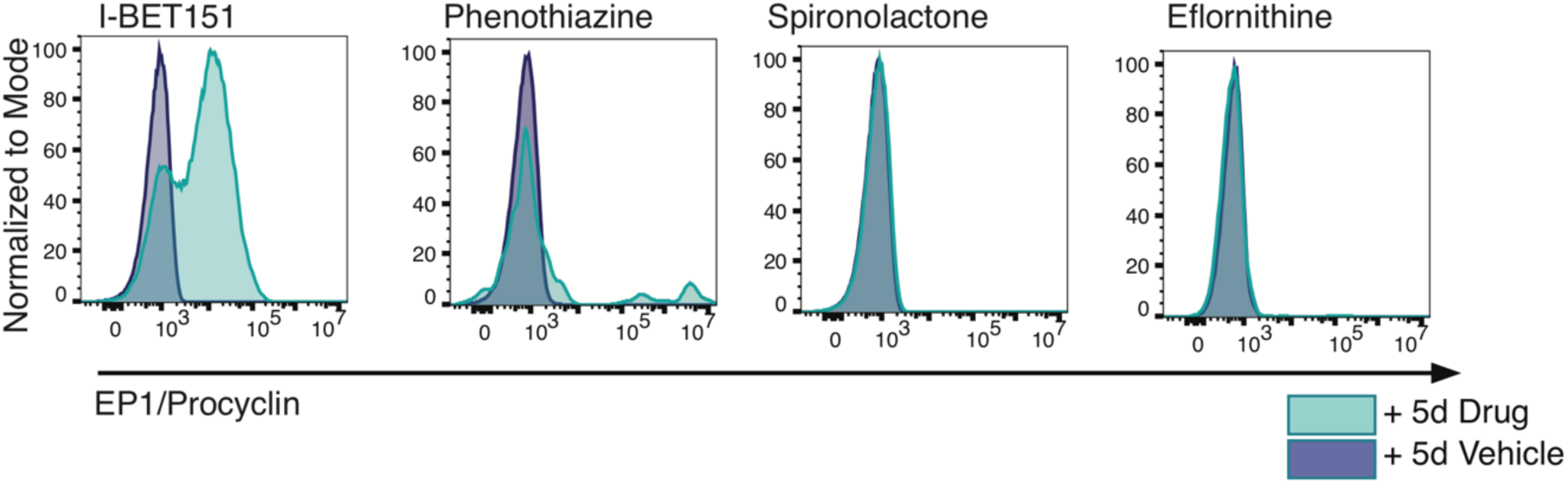
Treatment with phenothiazine, spironolactone and eflornithine does not result in detectable levels of procyclin protein on the cell surface. Flow cytometry plots of bloodstream parasites treated for 5 days with the indicated drug or vehicle control and stained with fluorescent anti-procyclin antibodies. The bromodomain inhibitor I-BET151 was used as a positive control.

**Figure 5.**
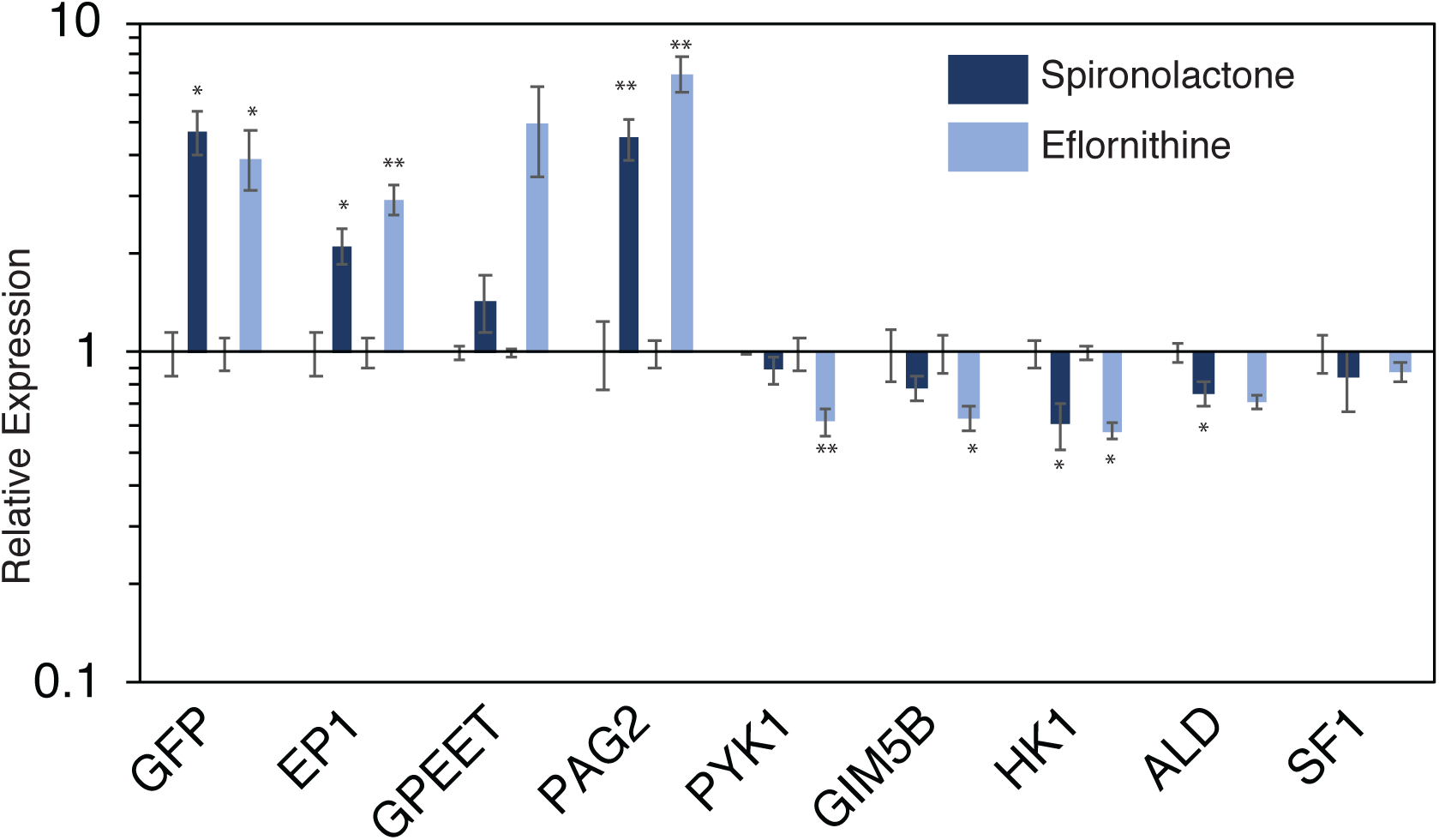
Treatment with spironolactone and eflornithine changes expression levels for a subset of differentiation associated genes. Q-PCR experiment showing relative expression levels for indicated genes in parasites treated for 3 days with 33µM spironolactone, eflornithine, or vehicle control. Error bars indicate standard error of 3 biological replicates. *, **, *** indicates p < 0.05, p < 0.01, and p < 0.001 respectively as measured by an unpaired, two-sided Student’s t-test. Note, relative expression is plotted on a log scale.

### The expression for a number of transcripts associated with differentiation is altered following treatment with spironolactone and eflornithine

Treatment with either eflornithine and spironolactone increased *EP1/GFP* expression in our bloodstream reporter cells, and this made us wonder whether other transcripts associated with differentiation show altered expression levels in drug treated parasites. While phenothiazine also increased *EP1/GFP* expression, we worried that this effect might be a non-specific stress response, and accordingly did not test phenothiazine’s effect on other differentiation-related transcripts. We treated parasites with either eflornithine or spironolactone for three days, isolated RNA, and performed qPCR to ask whether expression levels were altered for a subset of genes whose expression changes during differentiation to procyclic forms (30). In addition to an increase in *EP1* and *GFP* transcript levels for in drug treated cells, we also observed an increase in transcript levels for Procyclic Associated Gene 2 (*PAG2, Tb927.10.10220*), which is found just downstream of *EP1* in the same Polycistronic Transcription Unit (PTU) and is known to have increased expression during differentiation (Fig 5). Another surface protein expressed in early procyclic parasites (*GPEET, Tb927.6.510*) showed an increase in eflornithine treated parasites, but this increase was not statistically significant. Because differentiating parasites must alter their metabolism to adapt to the insect midgut, we also checked expression levels for a number of genes associated with these changes in metabolism. Both hexokinase (*HK1, Tb927.10.2010*) and glycosomal membrane protein (*GIM5B, Tb927.9.11600*) show an initial dip in expression levels following initiation of differentiation and then a slow increase in expression, while glycosomal fructose biphosphate aldolase (*ALD, Tb927.10.5620*) and pyruvate kinase (*PYK1, Tb927.10.14140*) show a monotonic decrease in expression following initiation of differentiation (30). We observed a subtle but statistically significant decrease in *PYK1, GIM5B*, and *HK1* following treatment with eflornithine and a decrease in *HK1* and *ALD* following treatment with spironolactone. No changes were observed in splicing factor 1 (*SF1, Tb927.10.9400*), which we used as a negative control as it has not previously been associated with changes in expression for differentiating parasites (Fig 5). It should be emphasized that changes in expression levels for differentiation-associated genes in our drug treated parasites are much lower than those observed in differentiating cells. Although the expression changes are muted compared to those in differentiating cells, it is interesting that small changes in *EP1* expression in eflornithine and spironolactone treated parasites are accompanied by other small changes in expression levels for a subset of genes known to be associated with differentiation. We conclude that the effect of both eflornithine and spironolactone is not solely confined to genes associated with changes in the proteins that make up the parasite surface, and may additionally initiate multiple components of the differentiation program.

### Identification of *in vitro* inhibitors for *T brucei* growth

Because our screen used a flow cytometer that can be programmed to analyze a fixed volume of cells, we were easily able to identify drugs that slowed trypanosome growth after 3 days of treatment by simply counting the number of analyzed cells that fell within a live cell gate. We binned each drug into four categories: highly inhibitory, inhibitory, slightly inhibitory, and not inhibitory (Table 1). Our DMSO vehicle treated cells averaged roughly 35,000 cells in the live gate at the end of the treatment. 35 drugs were classified as Highly Inhibitory because they resulted in 35 cells or less in the live gate. 102 drugs with 36-350 cells in the live gate were classified as Inhibitory, 17 drugs with 351-3,500 live gate cells were classified as slightly inhibitory, and any samples with greater than 3500 cells were classified as not inhibitory (the remaining 1,431 samples, Table 1). For the most highly inhibited category of drugs, we searched the literature to see if they had previously been characterized as anti-trypanosomal. Berberine, ipecac syrup, disulfiram, methylene blue hydrate, promethazine hydrochloride, perphenazine, promazine hydrochloride, and triflupromazine have all previously been reported to inhibit growth in *T. brucei* and our results agree with those previously published reports (31–35) (S2 Table, S3 Table). Several other drugs, including nitrofurazone, aminacrine, clemastine fumarate, bepridil hydrochloride, amiodarone hydrochloride, prochlorperazine dimaleate, protriptyline hydrochloride, fluoxetine hydrochloride, apomorphine hydrochloride, and proadifen hydrochloride have been shown to inhibit growth in *T. cruzi* (33,36–44). Finally, cyproheptadine hydrochloride was shown to inhibit growth in *T. evansi* (S2 Table) (45). We also identified a number of drugs that inhibit *T. brucei* growth *in vitro* that, to our knowledge, have not previously been studied in this context (S2 Table), including antimalarials, antibacterials, antiseptics, anesthetics, antihypertensives, antiemetics, antidepressants, vitamins, and a vasodilator. In addition to these highly inhibitory drugs, a further 119 drugs were shown to have a 10-100 fold inhibitory effect on parasite growth, including pentamidine, a well-established trypanosomiasis drug (S4 Table, S5 Table). In addition to assaying these drugs for their effect on trypanosome growth, we also measured percent survival after treatment for 24h with 16µM drug for two different cell lines: murine Raw264.7 macrophages and human C32 melanoma lines (S3-5 Tables). These additional data can be used to help prioritize the drugs with the highest mammalian cell line survival rate. We hope that this list can be used as a resource for those in the trypanosomiasis drug development field.

**Table 1.**
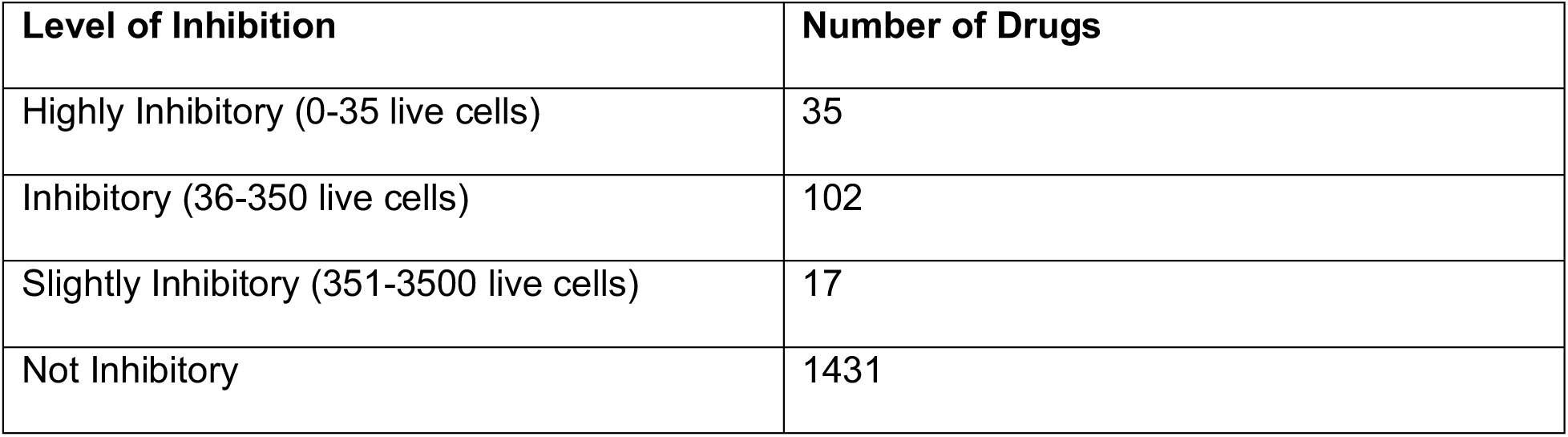
Summary of drugs that cause growth inhibition or cell death.

| Level of Inhibition | Number of Drugs |
| --- | --- |
| Highly Inhibitory (0-35 live cells) | 35 |
| Inhibitory (36-350 live cells) | 102 |
| Slightly Inhibitory (351-3500 live cells) | 17 |
| Not Inhibitory | 1431 |

### Confirmation that drugs identified in the screen inhibit trypanosome growth

To confirm that the drugs identified in the screen did indeed inhibit parasite growth, we ordered fresh stocks of six drugs from Sigma-Aldrich and measured growth inhibition. We chose three drugs from our most highly inhibitory category: flunarizine hydrochloride, aprepitant, and clemastine fumarate. Clemastine fumarate is an antihistaminic drug that has been shown to inhibit growth in *T. cruzi* (37). Aprepitant is an antiemetic and flunarizine hydrochloride is a vasodilator. To our knowledge neither of these latter two have been studied for their effects on kinetoplastids. In addition to these three highly inhibitory compounds, we chose one drug from our inhibitory category (triprolidine) and two drugs from our slightly inhibitory category (phenothiaizine and bufexamac). Neither triprolidine nor bufexamac have been shown to inhibit trypanosome growth, while phenothiazine has previously been shown to be strongly inhibitory (24,25). To assess the effect of each of these six drugs on trypanosome growth, we adopted a flow cytometry assay using the nucleic acid stain Sytox Orange, which positively stains dying and dead cells with compromised membranes. Parasites were plated at 100,000 cells/ml, and an aliquot of parasites was stained with Sytox Orange after 24h of growth. A fixed volume of cells was then analyzed by flow cytometry and the number of live cells was determined by gating on Sytox Orange negative populations whose size (measured by forward scatter) is consistent with healthy parasites (Fig 6A). Whereas healthy cells generally do not stain positively with Sytox Orange (Fig 6A, left), those treated with an inhibitory drug do show populations of positively staining Sytox Orange parasites as well as populations of smaller particles that represent dead cells and debris (Fig 6A, right). Based on the number of cells in the live gate, the remaining unstained parasites were diluted back down to 100,000 parasites/ml. The growth curves in Fig 6B show that parasites treated with five of the six drugs we tested had a severe growth inhibited phenotype, while treatment with triprolidine had no appreciable effect on growth (data not shown). Survival data for mammalian cell lines treated for 24h at 33µM for all verified *EP1/GFP* and growth inhibition drugs is provided in S6 Table.

**Figure 6.**
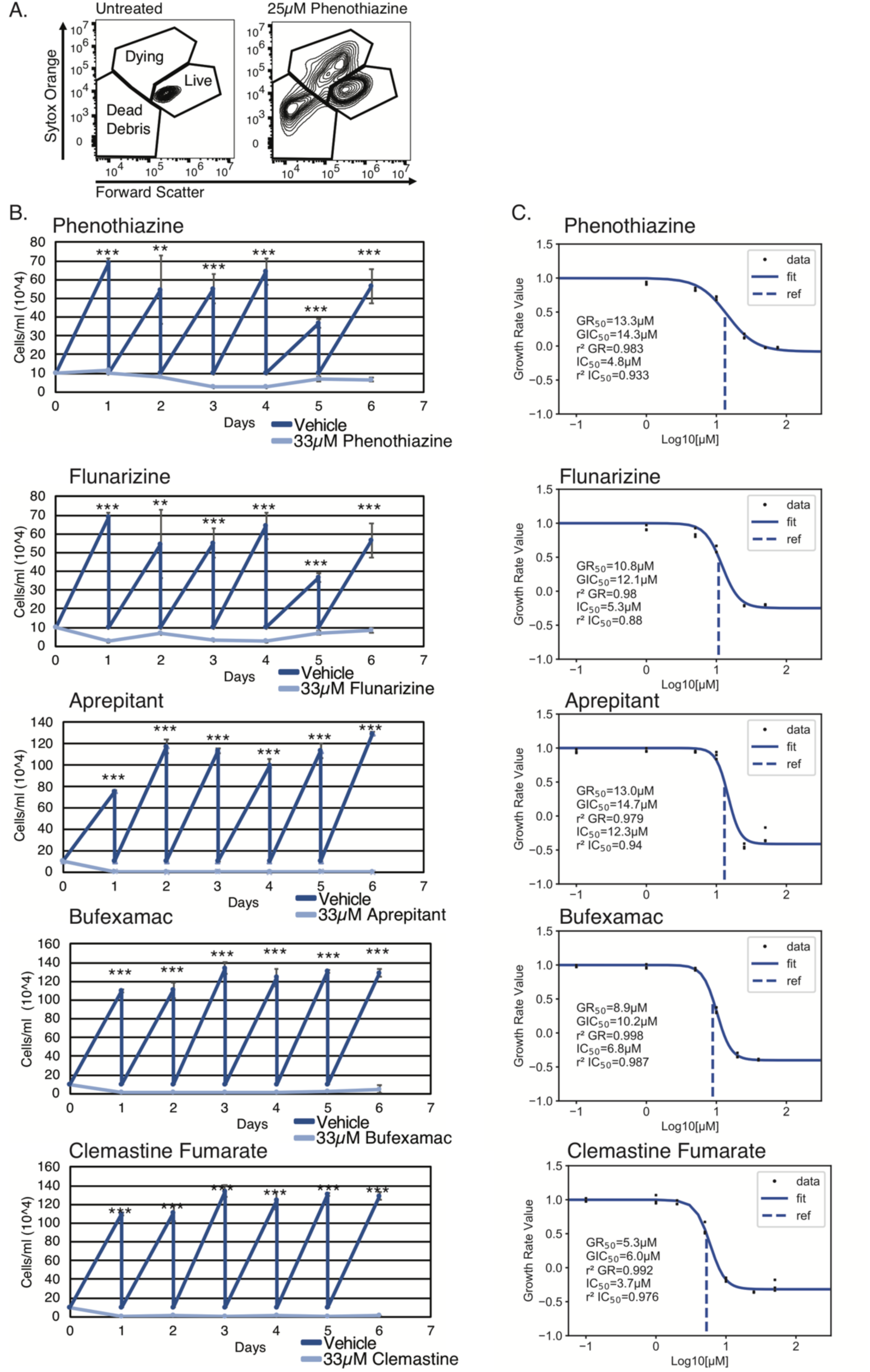
Confirmation of growth inhibition for a subset of drugs identified in the drug screen. (A) Flow cytometry plot showing forward scatter vs Sytox Orange staining to illustrate how live cells were gated. (Left) untreated parasites, (Right) parasites treated with 25µM phenothiazine. (B) Growth curves of parasites treated with the indicated drugs or vehicle control over a period of 6 days. Dark blue, vehicle control treated cells, Light blue, drug treated cells. Error bars indicate standard deviation of 3 biological replicates. *, **, *** indicates p < 0.05, p < 0.01, and p < 0.001 respectively as measured by an unpaired two-sided Student’s T test. (C) Percent growth inhibition over a range of concentrations for the indicated drug. Data were fitted and indicated values calculated using the GRMetrics R package. GR_50_, the concentration at which the effect reaches a growth rate (GR) value of 0.5 based on interpolation of the fitted curve (dashed lines). GIC_50_, the drug concentration at half-maximal effect for calculated growth rate. r^2^ GR, the coefficient of determination for how well the GR curve fits to the data points. IC_50_, the concentration at which relative cell count = 0.5. r^2^ IC_50_, the coefficient of determination for how well the traditional curve fits to the data points. See methods for more details.

We went on to estimate the IC_50_ values for each of the six drugs by growing parasites for 48h in a range of concentrations. Five of the six drugs showed IC_50_ values between 3.7 and 12.3µM (Fig 6C). Consistent with our observations on cell growth, the IC_50_ value measured for triprolidine was very high (S1 Fig) and the r^2^ value for this drug indicates a relatively poor fit for the data. Therefore, it should be noted that the screen can produce some false positives, and promising candidates need to be verified with fresh stocks prior to further study. Interestingly, we did not observe consistently higher IC_50_ values for drugs in the slightly inhibitory (phenothiazine, bufexamac) category when compared to those in the highly inhibitory category (aprepitant, clemastine fumarate, flunarizine). We conclude that compounds with only a 15-25- fold reduction in cell counts as measured in our screen can still have highly inhibitory effects on growth.

## Discussion

The work presented here provides proof of concept for a molecular tool that can be used to track the molecular events that initiate surface remodeling during trypanosome differentiation from the bloodstream form to the procyclic form, an event that is critical to life cycle progression. We first showed that transcription of *EP1/GFP* in our reporter line behaves as expected in bloodstream, procyclic and differentiating cells. One weakness of our reporter is that it was used in a monomorph strain rather than a pleomorph strain. While pleomorphs can be treated *in vitro* to closely mimic developmental transitions from bloodstream to procyclic parasites via the intermediate stumpy stage that is observed in the wild, monomorph lines are incapable of stumpy formation, making them a less physiological model. With this in mind, we created a pleomorphic *EP1/GFP* reporter strain and piloted it for use in flow cytometric screens. Unfortunately, *EP1/GFP* expression in these pleomorphic lines is much more variable than in the monomorph lines, and tends to increase with increasing parasite density (data not shown), which is consistent with preadaptation of stumpy parasites for transition to the insect vector (23). Therefore, we decided to use monomorph lines for screening purposes, and for any drugs that produced procyclin protein on the surface, these drugs could be separately tested in pleomorph lines using anti-procyclin antibodies.

This study demonstrates that our developmental reporter line can be used to conduct high-throughput flow-cytometry based small molecule drug screens. Importantly, our screening pipeline was developed such that growth inhibition can be tracked at the same time as the initiation of transcription for our gene of interest. This allowed us to quickly identify drugs that might cause a non-specific increase in *EP1/GFP* transcription, perhaps due to a stress response induced by the drug. It also allowed us to identify compounds that inhibited trypanosome growth, which can be used as a starting point for alternate trypanosomiasis therapeutics. We purposely used U.S. and foreign approved compounds because their toxicity is better characterized than those of new compounds, which might streamline drug development should promising candidates be found. However, there is no reason that other drug libraries could not be screened using the same methods described here.

We were able to identify three compounds that increased transcription of *EP1/GFP* after 3 days of treatment at 33µM. Phenothiazine’s effect on *EP1/GFP* transcription may well be non-specific, as the drug is a potent growth inhibitor, although many other drugs that inhibited growth did not have this effect. A previous study seeking to identify compounds that increase transcription of *EP1* in bloodstream cells resulted in 28 hits that all appeared to have some effect on cell growth (46). However, spironolactone and eflornithine did not severely inhibit trypanosome growth at 33µM, and showed reproducibly increased *EP1/GFP* transcription. Treatment with both spironolactone and eflornithine did not affect expression of *EP1* alone, as we found effects on several other differentiation-associated genes. This phenotype was also previously observed for I-BET151, although its effects on differentiation associated genes were of greater magnitude (17). In future, it would be interesting to test whether artificially increasing *EP1* levels via overexpression of the gene is sufficient to induce expression changes in other differentiation-associated genes.

While eflornithine had previously been reported to induce a stumpy-like, but developmentally incompetent, transition in monomorphic cells (22), the effect of spironolactone on *T. brucei* physiology has not been studied. In mammalian systems, spironolactone is an anti-androgen drug that non-selectively antagonizes mineralcorticoid receptors, including both progesterone and androgen receptors. It is used as a hypertensive agent to decrease morbidity and mortality of heart failure (47). To our knowledge, no proteins with similarity to mineralocorticoid receptors have been identified in *T. brucei*. We used BLAST to interrogate various kinetoplastid genomes with both the DNA-binding domain of the human glucocorticoid receptor and the ligand-binding domain of the mineralocorticoid receptor. While no hits were uncovered in *T. brucei*, we did obtain a few hits for genes in *T. cruzi* and *Leishmania panamensis* using the DNA-binding domain. All of the identified genes in *T. cruzi* (Tc00.1047053504039.110, Tc00.1047053506769.55, and Tc00.1047053510555.21) with an E value of less than 0.01 were for hypothetical proteins and ranged from 26%-33% identities. All three of them were also shown to be transcriptionally downregulated after infection in amastigotes and upregulated in epimastigotes (48). For BLAST with the ligand-binding domain of the mineralocorticoid receptor, we only obtained two hits with an E value of less than 0.05, one in *Perkinsela* (XU18_4875) and the other in *Leishmania Mexicana* (LMXM_29_2680). Both genes code for hypothetical proteins and had a 33-34% identity with the query. Future studies could potentially uncover the target of spironolactone in *T. brucei* by overexpressing a library of *T. brucei* ORFs and assaying for a decrease in *GFP* expression in spironolactone treated *EP1/GFP* reporter cells. These types of overexpression suppressor screens have previously been used successfully to identify drug targets in yeast and other organisms (49). With the gene target identified, genetic studies could uncover the role of this target (if any) in the initiation of transcription of *EP1* following the appropriate developmental signals in bloodstream form cells.

In addition to their effect on *EP1/GFP* transcript levels, both spironolactone and eflornithine affect expression levels of other genes known to be associated with differentiation (Fig 5). While we identified compounds that increased the transcription of *EP1/GFP*, we failed to identify any that produced procyclin protein on the surface of parasites, underscoring the already established paradigm that additional developmental steps may be required to remodel the parasite surface following transcription of the relevant RNAs. Given that the bromodomain inhibitor I-BET151 has previously been shown to increase both RNA and protein levels of procyclin (17), a more targeted screen using a library of epigenetic inhibitors might uncover more compounds that lead to this phenotype in the *EP1/GFP* reporter line.

Our screen uncovered 154 compounds that were slightly to highly inhibitory for cell growth. We were able to confirm a small subset of these by ordering fresh drugs from Sigma-Aldrich and performing growth and IC_50_ assays. 8 of the most highly inhibitory drugs had previously been shown to have effects on *T. brucei* growth, 1 had previously been shown to affect growth in *T. evansi*, and 10 had been shown to inhibit growth in *T. cruzi*. To our knowledge, the remaining 16 compounds that were in the highest category for growth inhibition have not previously been reported to affect *T. brucei* growth. We hope that our toxicity data can help to prioritize which of these drugs might be most promising as novel therapeutics. It is worth noting that pentamidine, a trypanosomiasis drug currently in active use, was sorted into the moderately inhibitory category, implying that the 102 compounds in this category could have potential as trypanosomiasis therapeutics. We were able to confirm the growth inhibitory effects of a small subset of drugs across the inhibitory categories by ordering fresh drugs from Sigma-Aldrich and performing growth and IC_50_ assays. One of the compounds we tested (triprolidine) was a false positive, for which we don’t have a good explanation. It is possible that evaporation of the drug plate resulted in some compounds being tested at higher than the intended concentrations, which would be consistent with the fact that we did find growth inhibitory effects of triprolidine at high concentrations in the IC_50_ assay. That said, we were able to confirm that the majority of compounds we tested had growth inhibitory effects consistent with the drug screen data. It is also worth noting that our screen likely contained some false negatives. Among other reasons, the hygromycin resistance drug used to select for the *EP1/GFP* reporter cells may have inactivated some of the drugs in the library due to phosphorylation.

While we focused on generating alternate trypanosomiasis therapeutics by inducing life cycle progression in *T. brucei*, this strategy could likewise be applied to other causative agents for neglected disease. Indeed, this strategy is actively being pursued for treatment of malaria, toxoplasmosis, and schistosomiasis (50–52). We think our reporter strain (and other developmental reporter strains such as the Pad1-GFP strain that tracks stumpy formation (53)) offer a powerful tool to perform genetic screens for developmental phenotypes. With the increasing availability of knockdown and overexpression libraries in *T. brucei*, there is great potential to uncover new life cycle-specific biological roles for the numerous proteins in *T. brucei* that are annotated as hypothetical. The combination of fluorescent reporter strains with high-throughput flow cytometry assays facilitates screening strategies that are both less labor-intensive and time-intensive than previous methods.

## Acknowledgements

We thank the members of the Harvey Mudd College Department of Biology for helpful discussions. We thank Monica Mugnier for helpful comments on the manuscript.

## Supplemental Figure Legend

**S1 Figure.**
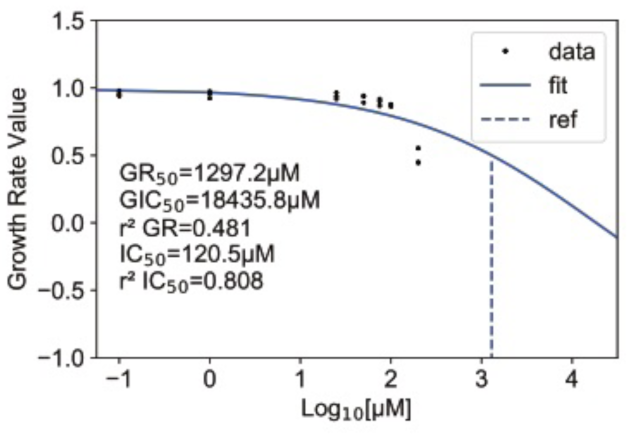
Triprolidine does not affect *T. brucei* growth at low concentrations. Percent growth inhibition over a range of concentrations for the indicated drug. Data were fitted and indicated values calculated using the GRMetrics R package. GR_50_, the concentration at which the effect reaches a growth rate (GR) value of 0.5 based on interpolation of the fitted curve (dashed lines). GIC_50_, the drug concentration at half-maximal effect for calculated growth rate. r^2^ GR, the coefficient of determination for how well the GR curve fits to the data points. IC_50_, the concentration at which relative cell count = 0.5. r^2^ IC_50_, the coefficient of determination for who well the traditional curve fits to the data points.

**S1 Table. Drugs that increase expression of *EP1/GFP***

**S2 Table. Information on the drugs identified as most highly inhibitory for *T. brucei* growth.**

**S3 Table. Drugs identified as most highly inhibitory for *T. brucei* growth. S4 Table. Drugs identified as most inhibitory for *T. brucei* growth.**

**S5 Table. Drugs identified as slightly inhibitory for *T. brucei* growth.**

**S6 Table. Toxicity values for drugs that verified to inhibit *T. brucei* growth.**

