## Supplementary figures and images for "Identification of clinically approved small molecules that inhibit growth and promote surface remodeling in the African trypanosome"

### Supplemental_Figure_1

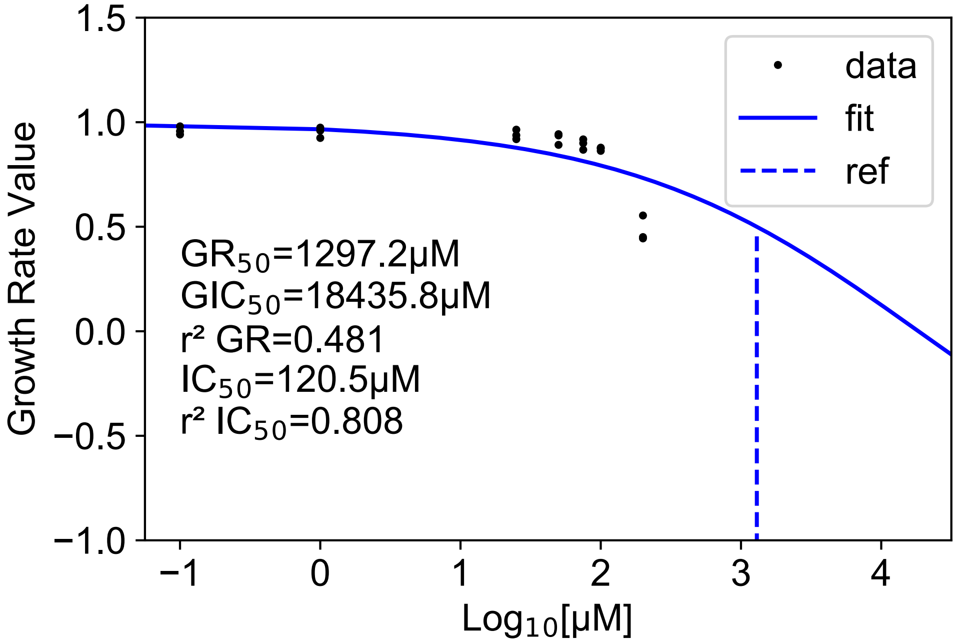
